# Small Molecule Activators of Protein Phosphatase 2A Exert Global Stabilising Effects on the Scaffold PR65

**DOI:** 10.1101/2025.07.24.666388

**Authors:** Mohsin M. Naqvi, Maria Zacharopoulou, Satyaki Saha, Anupam Banerjee, Zeynep S. Yilmaz, Vanda Sunderlikova, Chris M. Johnson, Janet R. Kumita, Shang-Hua Yang, Reuven Gordon, Michael Ohlmeyer, Sander Tans, Mert Gur, Ivet Bahar, Laura S. Itzhaki

## Abstract

Protein phosphatase 2A (PP2A), an important therapeutic target, comprises a scaffold subunit PR65 composed of 15 HEAT (Huntingtin/elongation/A-subunit/TOR1) repeats, a catalytic subunit, and one of many different regulatory subunits that enable binding to specific substrates. Recently, small molecule activators of PP2A (SMAPs) were identified, although their mechanisms of action have not been fully defined. Here we explore the interaction of PR65 with two SMAPs, ATUX-8385 and the non-functional DBK-776, using single-molecule optical tweezers, ensemble methods, and computational analysis. In the absence of SMAP, PR65 shows multiple unfolding and refolding transitions, and the force-extension profiles are very heterogeneous with evidence of misfolding. Similar heterogeneity has been observed for chemical-induced unfolding of tandem-repeat proteins like PR65, a consequence of the internal symmetry of the repeat architecture. In the presence of ATUX-8385, higher unfolding and refolding forces are observed globally, and there is less misfolding, suggesting that ATUX-8385 acts like a pharmacological chaperone. In contrast, DBK-766-binding induces higher unfolding forces for only a few repeats of PR65, suggestive of a more localised effect; moreover, subsequent stretch-relax cycles show that PR65 is irreversibly locked in the unfolded state. Docking and molecular dynamics simulations provide additional insights how SMAP binding modulates PR65 structure and function.

## Introduction

An intricate balance between kinase and phosphatase activities plays a vital role in signalling and protein homeostasis in the cell ^1,2^. Whereas many kinase inhibitors have been approved as treatments for human cancers, phosphatase inhibitors or activators have been less studied to date. Protein phosphatase 2A (PP2A), a serine/threonine phosphatase, belongs to a major class of enzymes regulating cell homeostasis by dephosphorylating key signaling molecules^3^. Its dysregulation has been associated with diseases such as cancer, neurodegenerative disorders (Alzheimer’s and Parkinson’s), and cardiovascular and pulmonary diseases, making PP2A an attractive target for therapeutic interventions^4,5^.

PP2A is a heterotrimer, composed of a scaffold (PR65 or subunit A), a catalytic subunit C, and a substrate-binding regulatory subunit B. A range of over 40 different B subunits, each specific for a distinct substrate or substrates, permits PP2A to control many different cellular signaling pathways. PR65 is a horseshoe-shaped tandem-repeat protein, composed of 15 HEAT (Huntingtin/elongation/A-subunit/TOR1) repeats of 39 amino acids each, whose sequence similarity is relatively low^2,6^. In the PP2A holoenzyme, the catalytic subunit binds to the C-terminal repeats 11-15 of PR65, and the regulatory subunit binds to repeats 1-10^3^.

Crystal structures of different PP2A heterotrimers and of uncomplexed PR65 alone, as well as studies of their conformational dynamics^7^, suggest that PR65 can adopt different conformations with varying degrees of compactness or extension and that PR65 needs to be highly flexible to accommodate the multitude of PP2A complexes necessary for diverse functionalities in the cell while maintaining its structural integrity. It has also been proposed that, rather than providing a rigid scaffold, PR65 is an elastic connector whose global motions allosterically coordinate cycles of catalysis of multiply phosphorylated substrates^7,8^. Indeed, the end-to-end distance of PR65 in different complexes differs by as much as 40 Å, indicating a remarkable scaffold flexibility^9^ amenable to modulation upon binding small molecules. Binding of small molecules could thus provide a mechanism for enhancing the holoenzyme activity.

Certain classes of tricyclic sulfonamides (narcoleptics) have been shown to bind and activate PP2A, thereby acting as potential therapeutic molecules^10–15^. A recent study indicates that one of these SMAPs (small molecule activators of PP2A), DT-061, stabilizes the heterotrimer upon binding to an inter-subunit pocket lined by all three PP2A subunits^16^. DT-061 was found to bind PR65 *in vitro*, with a dissociation constant of 235 nM, and the binding site was mapped to residues K194-L198 of PR65. Two other SMAPs, ATUX-8385 and its enantiomer ATUX-3364, have been studied for their effects on hepatoblastoma, a rare type of liver cancer. Both compounds effectively decreased the viability and proliferation of hepatoblastoma cells *in vitro*^*17*^.

In the present study, we explore the interaction of ATUX-8385 and DBK-766 (a non-functional SMAP) with PR65 using ensemble biophysical methods and examine their impact on protein folding using single-molecule optical tweezers^18^. The two SMAPs have distinct effects on hepatoblastoma H-1650 cell lines: Whereas ATUX-8385 induces cell death^19^, DBK-766 has no significant effect. Dissection of the unfolding and refolding pathways of PR65 under force in the absence and presence of these small molecules reveals a global ‘chaperoning’ effect exerted by the functional SMAP on PR65, reminiscent of the effect of a pharmacological chaperone on the folding behavior of the prion protein^20^. In the presence of the non-functional SMAP, on the other hand, PR65 exhibits smaller, more localized responses to uniaxial tension. We complemented the experimental analysis by docking simulations followed by molecular dynamics (MD), which revealed a novel site, designated as S3, for ATUX-8385 binding when PR65 is in its extended conformation. ATUX-8385 binding further stabilizes the extended form of PR65, as needed for facilitating the insertion and binding of the C and B subunits. The close proximity of this site to the structurally resolved DT-061-binding site (called S1 or S_exp_) suggests that the binding pose of the SMAP may be readjusted to enable optimal ternary contacts upon complexation of PR65 with subunits B and C. Notably, this new site does not accommodate stable DBK-766 binding, consistent with the “non-functional” nature of this particular SMAP.

## Results

### PR65 binds small molecules ATUX-8385 and DBK-766

The relatively low solubility of the small molecules did not allow the use of biophysical methods such as Isothermal Titration Calorimetry (ITC) to study binding to PR65. Consequently, alternative approaches were utilised. First, nano-Differential Scanning Fluorimetry (NanoDSF), a fluorescence-based label-free technique, was employed. NanoDSF measures the changes in the intrinsic tryptophan and tyrosine fluorescence of proteins upon thermal denaturation. A shift in the protein melting temperature (*T*_*m*_) can indicate either stabilisation or destabilisation of the protein structure globally or locally, induced by ligand binding. Thermal denaturation of 2 µM PR65 was measured in 10% DMSO, both in the absence and presence of 100 µM ATUX-8385 and 100 µM DBK-766. A increase in the melting temperature of PR65 was observed with both ATUX-8385 and DBK-766 (from *T*_*m_PR65*_ = 52.7 ± 0.1 °C to *T*_*m*_*_*_*PR65+ATUX*_ = 53.5 ± 0.1 °C, and *T*_*m_PR65+DBK*_ = 53.5 ± 0.1 °C), indicating binding and a small degree of stabilisation of the protein structure (**Fig. 1a-c**). The small shifts in *T*_*m*_ suggest dissociation constants in the micromolar range.

**Fig. 1.**
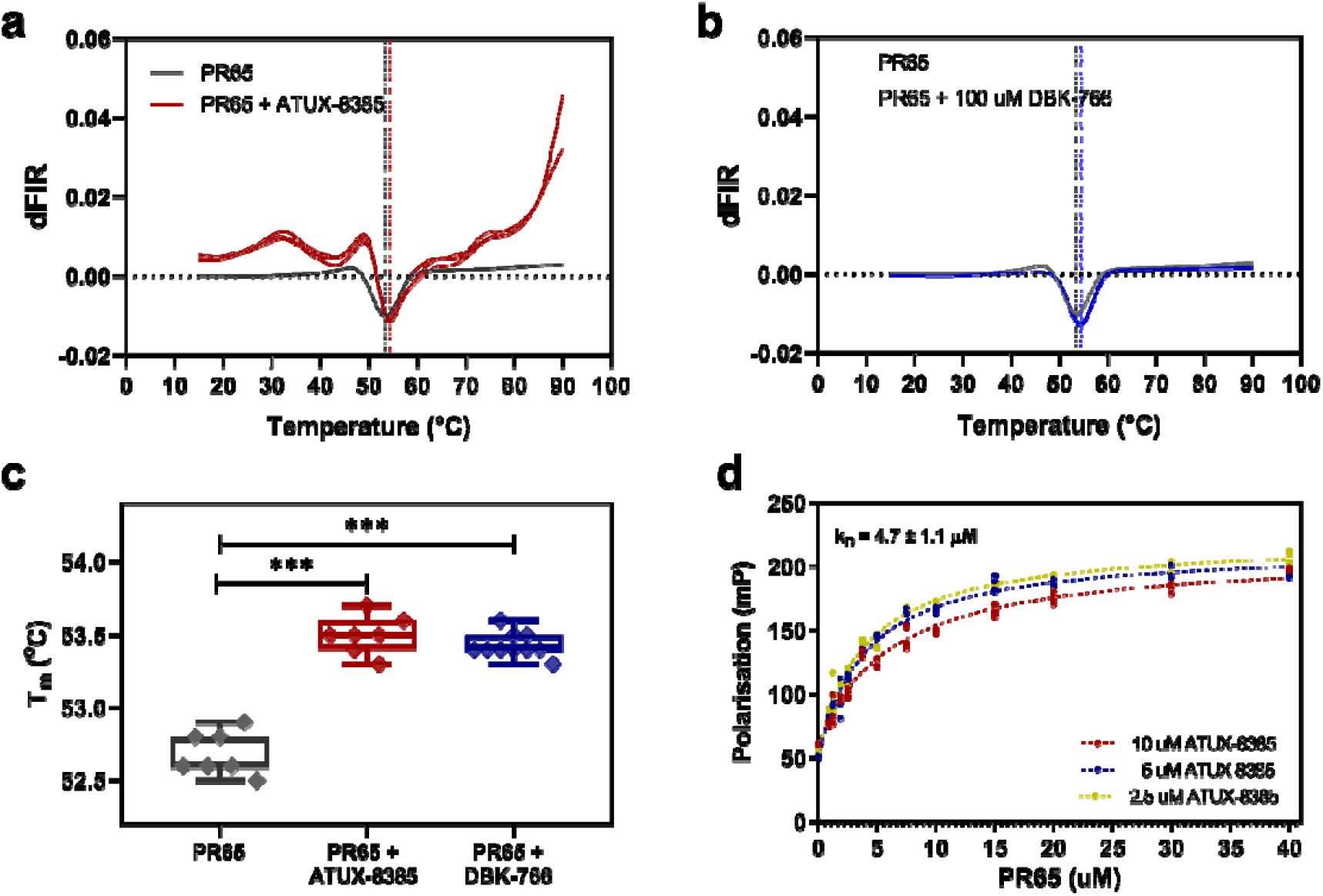
Small molecules ATUX-8385 and DBK-766 bind to PR65. **a)** NanoDSF traces of the thermal denaturation of PR65 in the absence (grey traces) and in the presence of ATUX-8385 (red traces). The data shown are the first derivative of the ratio of fluorescence intensity read at 350 nm over at 330 nm (dFIR (350 nm/330 nm)). The global minimum corresponds to the melting temperature of the protein, T_m_. A shift towards higher T_m_ values indicates an increase in stability induced upon SMAP binding (*T*_*m_PR65*_ = 52.7 ± 0.1 °C, *T*_*m*_*_*_*PR65+ATUX*_ = 53.5 ± 0.1 °C). **b)** NanoDSF traces of the thermal denaturation of PR65 in the absence (grey traces) and presence (blue traces) of DBK-766. Again, *T*_*m_PR65+DBK*_ = 53.5 ± 0.1 °C. **c)** Extracted T_m_ values from the NanoDSF traces indicate an upwards shift in the presence of ATUX-8385 and DBK-766 (p < 0.001 via ordinary one-way ANOVA with multiple comparisons). The NanoDSF experiments were performed with a Prometheus NanoDSF instrument (NanoTemper Technologies), 2 µM of PR65 in PBS, 2 mM DTT, incubated either with 10% DMSO, 100 µM ATUX-8385, or 100 M DBK-766, in a final concentration of 10% DMSO, and thermal denaturation was performed from 20 °C to 90 °C with a 1 °C/min rate. **d)** Fluorescence polarisation experiments show ATUX-8385 binding on PR65 with a dissociation constant in the low micromolar range. Shown are the fluorescence polarisation values (mP) for 2.5 µM, 5 µM, and 10 µM ATUX-8385 upon PR65 titration, N=3. One-site fitting of the data (see Methods) gives a *K*_D_ of 4.7 ± 1.1 µM.

To quantify the binding of ATUX-8385 to PR65, the fluorescent properties of ATUX-8385 were exploited. Upon excitation at 320 ± 10 nm, ATUX-8385 showed a maximum fluorescence intensity at 370 nm (**Fig. S1**). Fluorescence polarisation (FP) was employed to determine the dissociation constant. FP is based on the principle that the rotational motion of a fluorescent molecule affects the polarisation of the emitted light. Upon binding to PR65, the rotational motion of ATUX-8385 was reduced, resulting in higher polarisation values. PR65 was titrated into ATUX-8385 (10% DMSO) at 2.5 µM, 5 µM, and 10 µM (**Fig. 1d**), which gave a dissociation constant (*K*_D_) of 4.7 ± 1.1 µM. DBK-766 is not fluorescent (**Fig. S1**), precluding determination of its binding affinity by FP.

### Multiple (un)folding pathways of PR65 detected by optical tweezers

Single PR65 molecules were tethered between polystyrene beads held in dual optical traps^21^ via 600 bp DNA handles (**Fig. 2a**, see Methods)^22,23^. The constant velocity pulling experiments at 100 nm/s displayed a heterogeneous unfolding/refolding behaviour with hysteresis. Multiple intermediate states were observed during stretching curves with variations in extension, force, and number of transitions in consecutive pulls and in different molecules (**Fig. 2b-e**, *red curves, red arrows*, **Fig. S2a**). The series of distinct unfolding transitions with varying extensions indicate that PR65 unfolds via domains of different sizes and stabilities due to varying cooperativity between the repeats and/or helix motifs^6,24^. Similar heterogeneity was also observed during relaxation/refolding curves (**Fig. 2b-c**, in *blue, blue arrows*). Interestingly, PR65 also displayed misfolding behaviour^25^ marked by stretching traces that showed intermediate states that could not be fully unfolded even when high forces were applied (**Fig. 2d-e**, *middle and right panels*). Notably, these states were not observed in the first pulls (**Fig. S2b**) and occurred abruptly in any successive stretching cycle (**Fig. 2d** and **e**), suggesting that the behaviour was due to the incorrect folding of some subdomains that disrupted the folding of the entire protein.

**Fig. 2.**
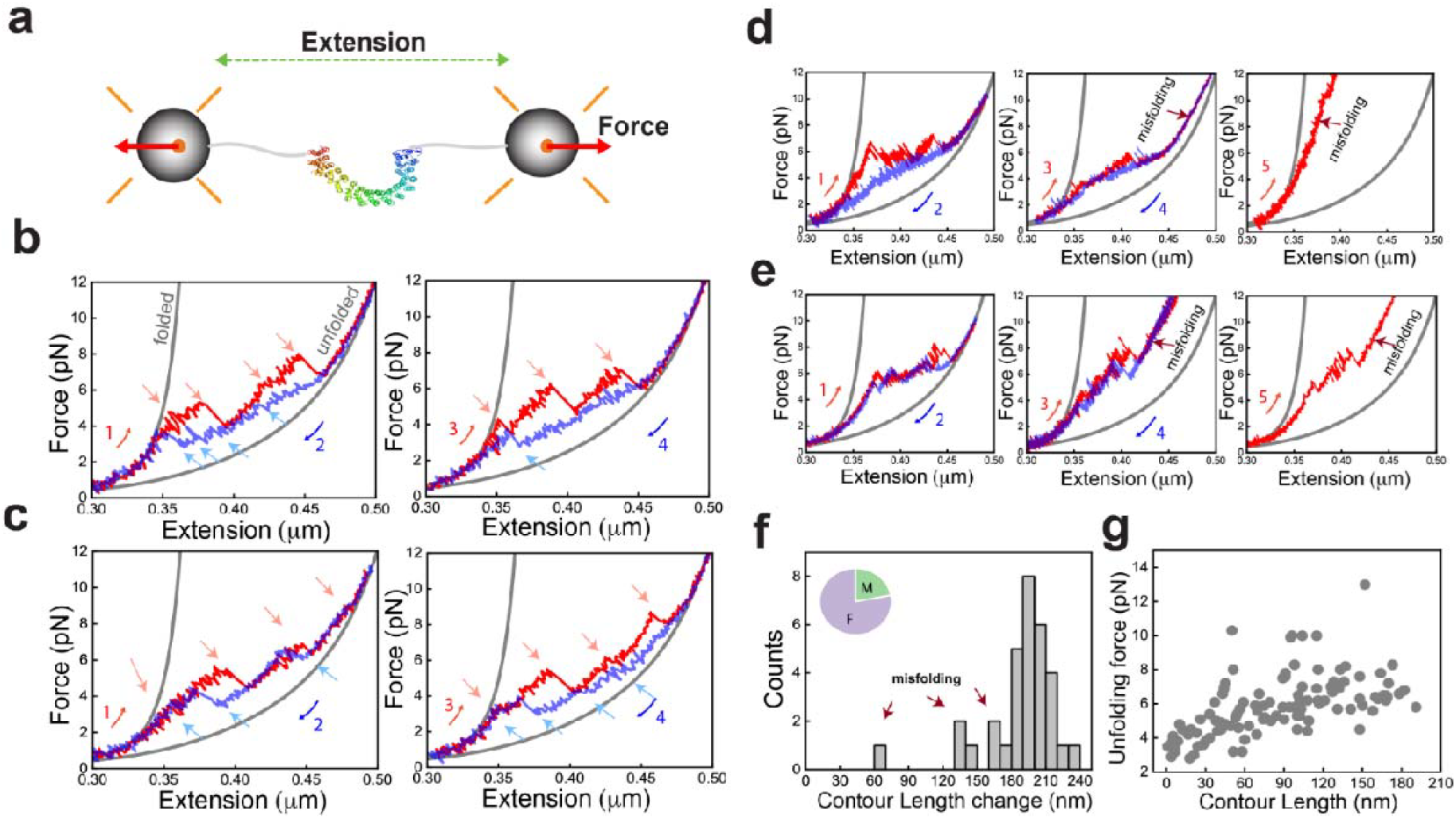
Single molecule (un)folding of PR65. **a)** Mechanical manipulation of single PR65 molecules using dual beam optical tweezers setup. **b)** Stretch (*red*) - relax (*blue*) cycles showing multiple intermediate states and heterogeneity in the lengths of unfolding (*red arrows*) /refolding (*blue arrows*) rips and number of intermediates in successive cycles. Numbers denote the order of the pull-relax cycles. *Gray* curves are WLC fitting. **c)** Example force extension traces of another PR65 molecule showing variability in the stretching and relaxation curves compared to molecule in **b. d-e)** Example force extension traces for molecules showing abrupt misfolding events in subsequent stretching curves after the first pull-relax cycle. **f)** Contour length change (ΔL_c_) distribution from folded to unfolded state (**b**) from WLC fitting of stretching curves (N_cycles_ = 32, N_molecules_ = 16). Inset shows percentage of stretching curves that displayed misfolding (M) and full folding (F). **g)** Unfolding force *vs* absolute contour length (L_c_) of each intermediate state observed during pulling (N_cycles_ = 32, N_molecules_ = 16).

We next fitted the folded and unfolded branches (**Fig. 2b**, *gray curves*) of the force-extension curves (FECs) to the extensible Worm-like chain (WLC) model^26^ (Methods). The distribution of contour length change of unfolding (ΔL_c_) displayed a major peak at 198 ± 1.4 nm matching with the expected length of 200 nm (**Fig. 2f**). The average force for complete unfolding of PR65 was determined to be ~ 7.2 pN at 100 nm/s pulling speed. The shorter lengths in the distribution correspond to the misfolded fraction (22% of the total stretching traces) that could not be fully unfolded even when high forces were applied (**Fig. 2f** *inset*).

We quantified the force and absolute contour length (L_c_) prior to each unfolding and refolding rip (**Fig. 2g;** see Methods) from those traces that showed no misfolding. The absence of clustering of data points indicates population of multiple intermediate states and a complex network of interactions between the helix-motifs. These intricate interactions result in the distinct unfolding transitions of domains with varying stabilities in each stretch-relax cycle. The heterogeneity in the unfolding/refolding forces and contour lengths of transitions in our single-molecule experiments is highly reminiscent of the behaviour observed for chemical-induced unfolding of PR65 in ensemble measurements and shown to arise from multiple pathways^6^. The presence of multiple pathways of similar energy for (un)folding has been reported by our group and others for several repeat proteins^27–32^. This feature reflects the high internal symmetry and is in striking contrast to globular proteins for which there are generally only single pathways accessible (for both chemical- and force-induced unfolding)^33,34^.

### ATUX-8385 binding globally stabilizes PR65 and prevents misfolding

To understand how functional SMAPs interact with PR65, we repeated the optical tweezers experiments at 100 nm/s in the presence of ATUX-8385 (**Figs. 3 and S3**). Interestingly, most PR65 molecules displayed higher unfolding forces compared to those in the absence of ATUX-8385, for each transition during stretching (compare **Figs. 2b-c** and **3b-d**); and importantly, the higher resistance to deformation was observed starting from the first pulls indicating a increase in the stability of the folded state in the presence of the functional SMAP (**Fig. 3b-c**, *red curve* and **Fig. S3b**). A subset of molecules showed stabilization of only a few subdomains of PR65, indicating a different binding mode (**Fig. 3d**). Similar increases in the refolding forces were also observed during relaxation, with variation in the hysteresis between stretch-relax cycles (**Fig. 3b-c,g and Fig. S3c**). The heterogeneity in the extensions, forces, and number of transitions within successive stretch-relax cycles and between different molecules was similar in the presence and absence of SMAP (**Figs. S2a** and **S3a**). Notably, and in striking contrast to the no-SMAP data, no misfolding was observed in the presence of SMAP, with ΔL_c_ values showing a single sharp peak at 196±3 nm indicating high population of the folded state (**Fig. 3e**).

**Fig. 3.**
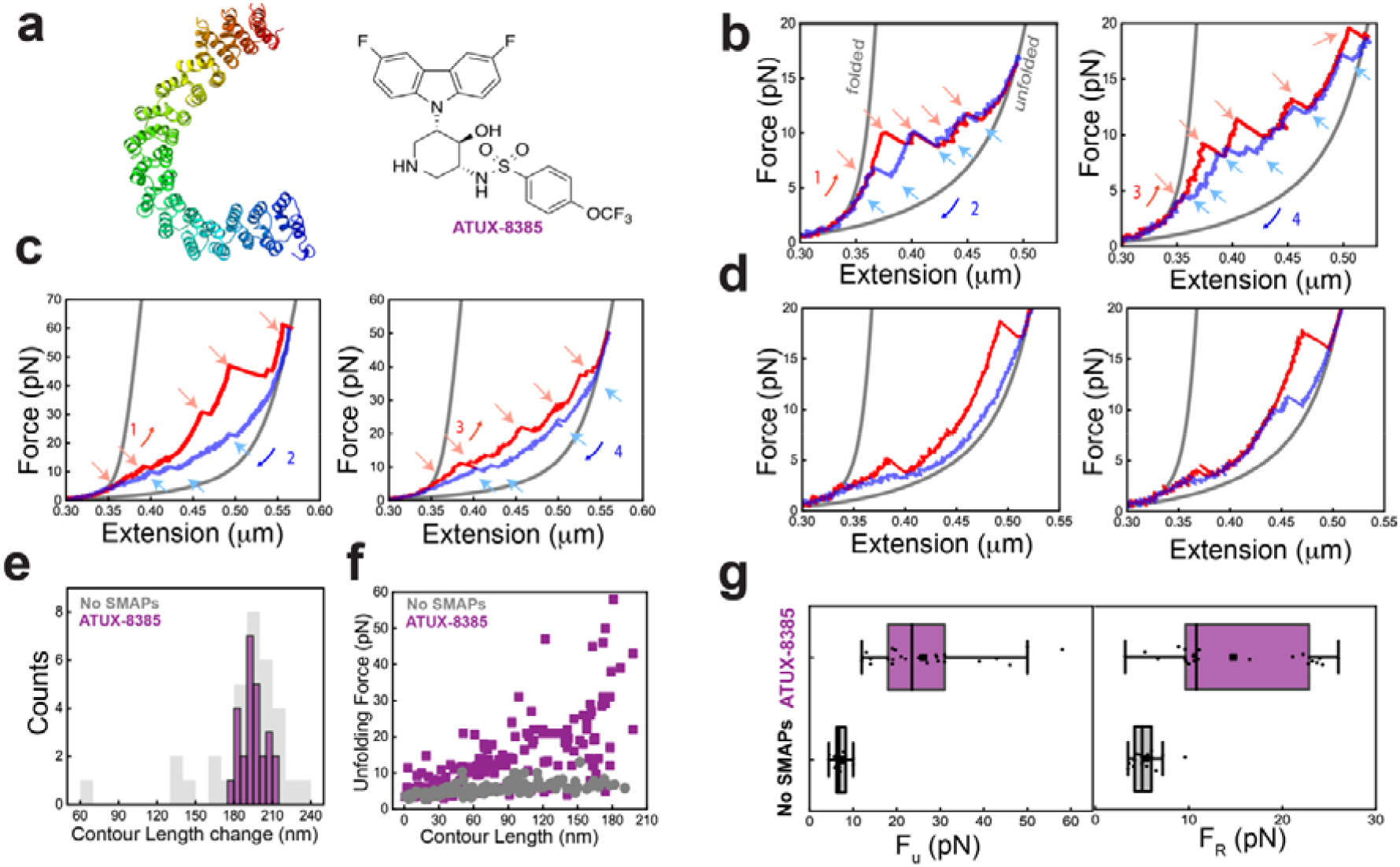
ATUX-8385 acts as chaperone in the folding of PR65. **a)** Structure of the tricyclic neuroleptic compound ATUX-8385. **b)** Force-extension curves of PR65 from pull-relax experiments at 100 nm/s, in the presence of 10 µM SMAP dissolved in < 2% DMSO in Tris-HCl buffer (see further details in Methods). Stretching (*red*) and relaxation (*blue*) curves from two consecutive cycles showing high unfolding and refolding forces of each rip. Gray curves are WLC fitting. **c)** Example force-extension traces from consecutive cycles from another PR65 molecule displaying heterogeneity in the unfolding and refolding pathways with larger hysteresis (*left panel*) compared to the molecule in **b**). **d)** Example force-extension traces from two separate PR65 molecules (*panels right and left*) showing stabilization of a few domains of PR65. **e)** ΔL_c_ distribution quantified (in *purple*) from all the stretching curves (N_cycles_ = 26, N_molecules_ = 16) in presence of ATUX-8385 showing a single peak corresponding to the folded PR65 in contrast to the misfolding fractions observed for the no-SMAP condition (in *gray*). **f)** Unfolding force versus absolute contour length (L_c_) of each intermediate state (*purple* with SMAP, *gray* without SMAP (from **Fig. 2g**) observed during pulling (N_cycles_ = 26, N_molecules_ = 16). Data showing ATUX-8385 binding stabilizes the entire PR65 folded state. **g)** Box plots showing distribution of maximum unfolding force (*left panel*) and refolding force (*right panel*) with (N = 26, 24, *purple*) and without (N=23, 19, *gray*) SMAP.

The large difference in the plots of unfolding force *versus* L_c_ in the presence *versus* absence of SMAP (**Fig. 3f** and *left panel* of **Fig. 3g**) suggests that SMAP binding globally stabilizes the folded state of PR65. This effect could arise from SMAP binding at multiple sites or binding at a single binding site having a long-range effect on the folded structure, increasing the thermodynamic stability and the barrier to unfolding. The interactions of the SMAP with PR65 also prevent its misfolding.

### Binding of non-functional SMAP, DBK-766, impedes the folding of PR65

When the unfolding experiments were performed in the presence of the non-functional SMAP, DBK-766, the data for the first pulls show that there are higher unfolding forces compared with the no-SMAP condition but for only a few repeats of PR65 as (**Fig. 4a-b**, *left panels*, **Fig. 4c**). This behaviour was also observed in the small subset of traces in presence of ATUX-8385 (**Fig. 3d**), indicating a limited mode of interactions in those cases. Interestingly, however, in subsequent stretch-relax cycles no significant unfolding/refolding transitions were observed **(Fig. 4a-b** *right panels*), and the PR65 molecule appeared to be locked in the unfolded state for multiple cycles before breaking. Since this locking behaviour was never observed for either ATUX-8385 or no-SMAP conditions, it suggests a distinct mode of interaction of DBK-776 with the unfolded state of PR65. When all runs are plotted (**Fig. 4d**), the ΔL_c_ distribution thus showed two distinct peaks for the folded and unfolded lengths (33% of traces). DBK-766 has a significantly different mode of interaction with both the folded and unfolded states of PR65 as compared with the functional ATUX-8385 SMAP **(Fig. 4c)**. These data suggest that DBK-766 binds weakly to the folded state but significantly stabilizes the unfolded state of PR65 hindering the folding of the protein.

**Fig. 4.**
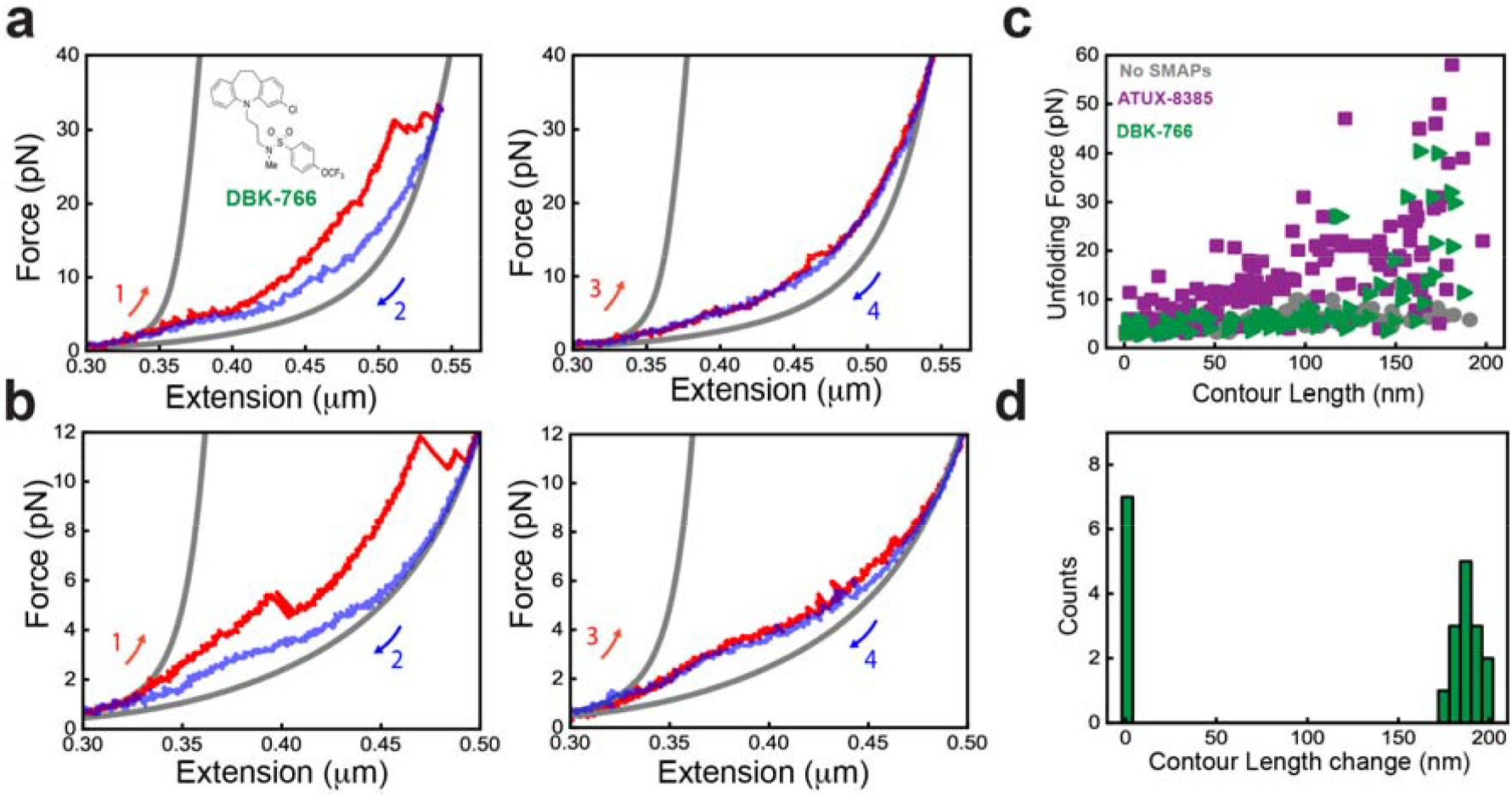
DBK-766 binding impedes the folding of PR65. **a-b)** Force extension curves of PR65 pull-relax experiments at 100 nm/s, in presence of 10 µM DBK-766 SMAP (*inset*) dissolved in < 2% DMSO in Tris-HCl buffer (see Methods). Stretching (*red*) curve (*left panel*) showing high unfolding force of a few repeats while no significant refolding jumps were observed during relaxation (*blue*). In the next pull-relax cycle (*right panel*) the molecule remained unfolded. Gray curves are WLC fitting. **c)** Unfolding force *vs* absolute contour length (L_c_) of each intermediate state observed during pulling (with DBK-766 in *green*: (N_cycles_ = 21, N_molecules_ = 10), with ATUX-8385 in *purple* (from **Fig. 3f**), without SMAP in *gray* (from **Fig. 1g**). Data showing weak stabilization by DBK-766 binding as compared to ATUX-8385 binding. **d)** ΔL_c_ distribution quantified from all the stretching curves (N_cycles_ = 21, N_molecules_ = 10) in presence of DBK-766 showing one peak corresponding to the folded PR65 and another peak at 0 nm corresponds to the pulls showing no significant unfolding transitions (**Fig. 4a-b** *right panels*).

### A second site (S3 or S_ATUX_), in proximity of the site (S_exp_) resolved for DT-061, shows high affinity for binding ATUX-8385

Both ATUX-8385 and DBK-766 are tricyclic sulfonamides, but the above results show that they exhibit different responses to uniaxial tension. Here we undertook a deeper examination of the potential binding sites of these two SMAPs to PR65 toward better understanding the molecular basis of their distinctive behaviour observed in force-extension experiments. Among the known SMAPs, only DT-061 complexed with PP2A trimer has been structurally resolved to date^16^: this SMAP sits at the interface of three subunits of PP2A. Previous label-free single-molecule experiments with nanoaperture optical tweezers combined with MD simulations gave first insights into the effects of ATUX-8385 binding on PR65 conformational behavior, and its optical scattering properties^35^. The results also pointed to the possibility of ATUX-8385 binding to a new site, designated hereafter as S3, alongside S2 mentioned above. Notably, S2 is the preferred site for binding to the compact form of PR65. It is consistent with hydroxyl radical footprinting experiments which identified the PR65 K194-L198 segment as a part of the putative SMAP-binding region for a different tricyclic molecule. On the other hand, by gradually increasing the docking simulation box size to 80 × 80 × 80 Å^3^ in the vicinity of the K194-L198 patch a broader diversity of binding sites is detected, with site S3 distinguished by its high propensity (see Methods for details). This site (also called S_ATUX_) neighbors the site S1 (or S_exp_) resolved by cryo-EM for DT-061-bound to PP2A, as illustrated in **Fig. 5a**. The figure shows the two alternative conformations of PR65, extended (*light green*, adopted as starting conformer in simulating ATUX-8385 (*red, space-filling*) binding), and compact (*dark green*, resolved by cryo-EM for the trimer complexed with DT-061) together with ATUX-8385 bound to S3 or S_ATUX_, as predicted by simulations. The extended PR65 conformation was used in docking simulations as it was the structure resolved and preferentially assumed in the absence of other subunits. **Fig. S5b and S6a** present further results from the docking simulations that identified S3. As shown in **Fig. 5b**, ATUX-8385 binds to a pocket lined by the inner helices of the HEAT repeats 3-5. **Fig. S6a** shows the binding pose from a different perspective.

**Fig. 5.**
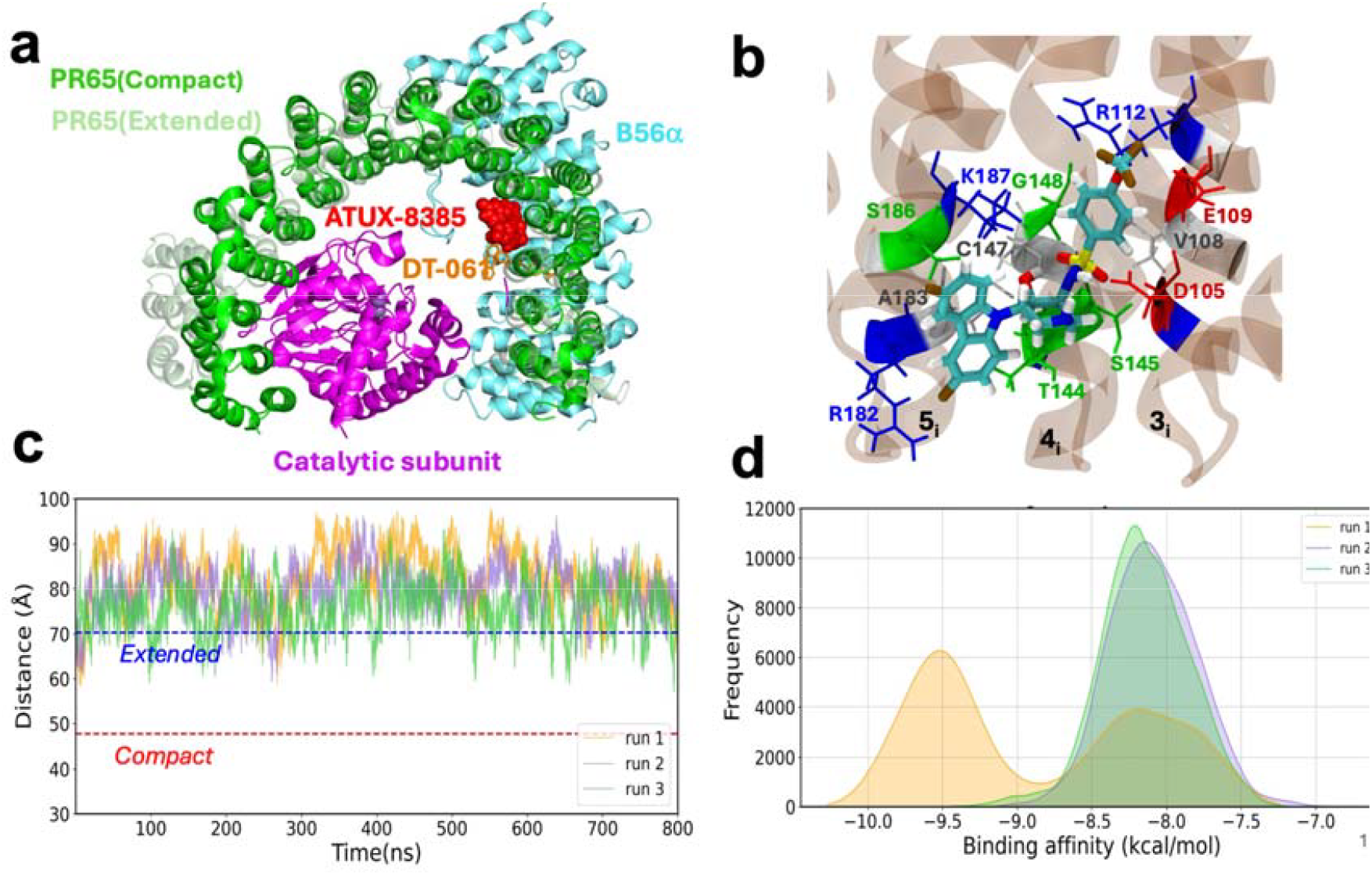
Results from docking and molecular simulations of ATUX-8385 binding to PR65. **a)** Comparison of the binding site and dynamics of SMAP DT-061 to PP2A (resolved by cryo-EM) and that of ATUX-8385 to PR65 (predicted by docking simulations). Simulations were performed for the extended form of PR65 (PDB: 1B3U^55^), while the PR65 in the DT-061-bound PP2A structure has a compact conformation. The two structures are optimally aligned to visualize the poses of the two SMAPs. **b)** Representative pose of ATUX-8385 at Site S_ATUX_ in the complex with PR65. ATUX-8385 is shown in *teal* licorice representation. The coordinating residues of PR65 from inner helices of repeats 3-5 (designated as 3_i_, 4_i_, and 5_i_) are shown and labeled. PR65 is rendered with a transparent brown background. **c)** Time evolution of PR65 end-to-end distance in the presence of ATUX-8385. Three independent MD runs of 800 ns each were performed. ATUX-8385 maintains its original binding pose, while PR65 maintains its extended conformation in all three runs **d)** Distribution of binding energies observed in MD simulations. The binding affinities of conformers visited during the three MD runs were evaluated using PRODIGY-LIG. Histograms corresponding to runs 1, 2, and 3 are shown by *orange, purple*, and *green*, respectively.

A series of MD simulations were conducted to evaluate the stability of ATUX-8385 bound to site S_2_ and the conformational dynamics of PR65 in the ATUX-8385-bound form. ATUX-8385 binding was observed to stabilize PR65 in an extended conformation, evaluated by measuring end-to-end distance between N29-F577 (**Fig. 5c** and **S6c**). As shown in **Fig. S6d**, ATUX-8385 maintains its original pose at S3 in two of the runs (with a root-mean-square deviation (RMSD) of < 5 Å with respect to the starting pose; whereas the RMSD increases to ~ 8.5 Å at around 350 ns in the 3^rd^ run. The sudden hike in RMSD originates from the flipping of ATUX-8385 at the same site S_ATUX_, rather than being displaced elsewhere. In parallel, the binding affinity of ATUX-8385 shows two peaks in the same run, at −8.2 kcal/mol and −9.5 kcal/mol (**Fig. 5d**), the latter corresponding to the flipped orientation. The other two runs show unimodal distribution of affinities with a peak around −8.2 kcal/mol, consistent with the positioning of the activator at the same site and same orientation during the course of these two simulations.

### DBK-766 preferentially binds an alternative site that does not allow for engaging in ternary interactions that promote the assembly of PP2A subunits

Similar simulations performed DBK-766 revealed a different behaviour. Binding of DBK-766 to S_ATUX_ was substantially weaker. Extensive MD simulations revealed its instability, as illustrated by the broad distribution of its RMSD in **Fig. 6a**, in comparison to that of ATUX-8385. In all three MD runs (of 800 ns each), DBK-766 RMSD with respect to its original pose exceeded 25 Å. **Fig. 6b** showcases one of such instances, where dislocation of DBK-766 from site S_ATUX_ is observed, followed by complete dissociation at 333 ns during MD run1. Since S_ATUX_ is not stable for DBK-766, the molecule begins to move around PR65, temporarily binding at various positions inside the repeats.

**Fig. 6.**
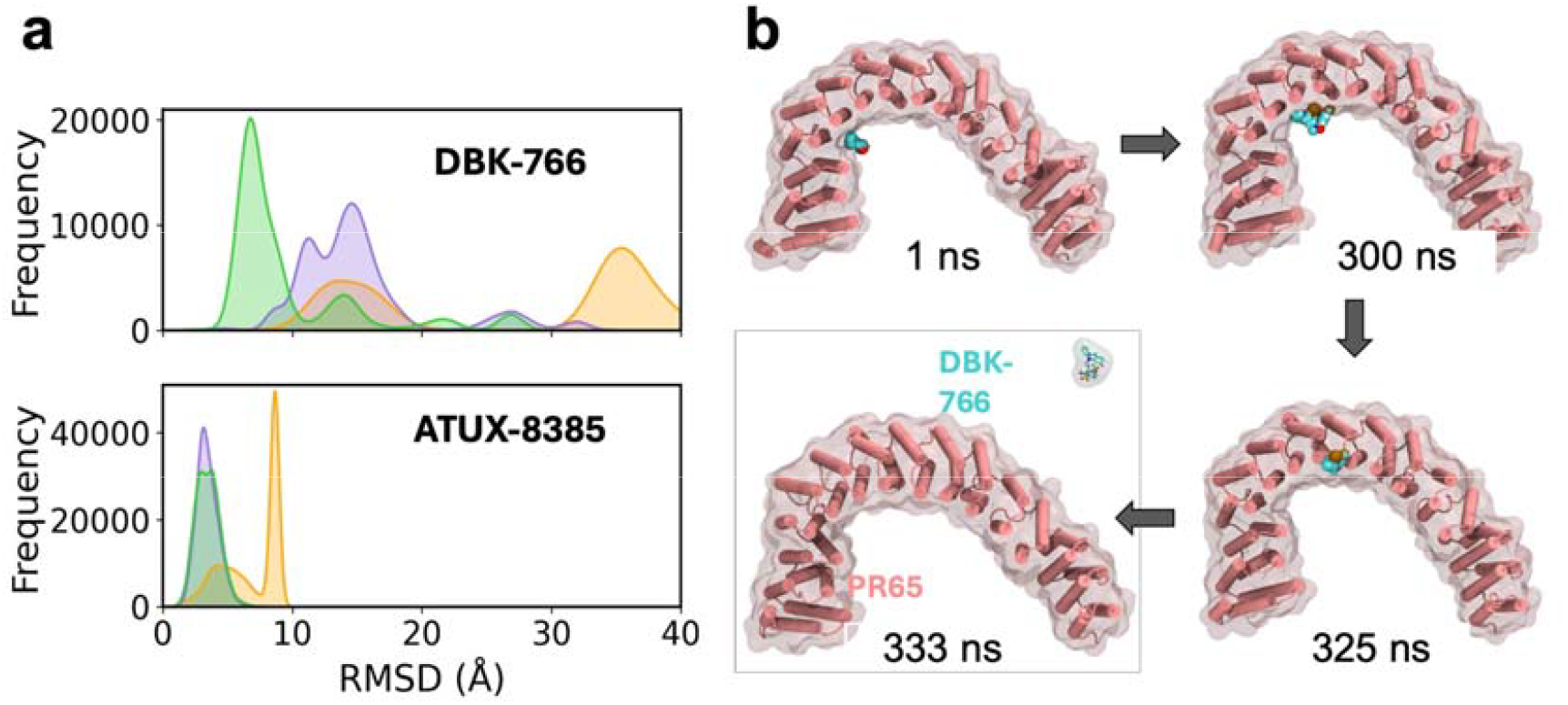
DBK-766 is not stable at site S_ATUX_. **a)** *(Top panel)* RMSD histogram of **DBK-766** from MD simulation snapshots shows higher RMSDs with respect to the initial pose, indicating instability and dissociation from site S_2_. *(Bottom panel)* In contrast, the RMSD histogram for **ATUX-8385** remains below 10 Å. MD runs 1, 2, and 3 are represented in *orange, purple*, and *green*, respectively. **b)** Four snapshots illustrating the gradual dislocation and dissociation of DBK-766 observed in MD run 1.

DBK-766 is a non-functional SMAP, but experimental evidence indicates that it is capable of binding to PR65, suggesting the existence of an alternative stable binding site that is not functional for PP2A holoenzyme formation/activation) for DBK-766. To find that, we further analyzed our docking simulations and identified another novel binding site, located along the outer (convex) surface of the HEAT repeats 7 and 8 (7_o_ & 8_o_) (**Fig. 7a**). MD runs conducted in triplicate starting from DBK-766 bound to that site showed that DBK-766 remained stably bound in two runs (runs 2 and 3), while a complete dissociation took place in run 1, as depicted by the SMAP RMSD (**Fig. 7b**). Excluding the latter from further analysis, we observed that DBK-766 also favoured the extended end-to-end distance for PR65 (**Fig. 7c**). Its binding affinity evaluated using *PRODIGY-LIG*^36^ ranged from −7.5 kcal/mol to −10 kcal/mol in both runs, similar to that of ATUX-8385 binding to S_2_ (**Fig. S7**). However, being on the exterior of the horseshoe-like PR65 structure, DBK-766 cannot interact with the other two subunits in favour of a more compact form conducive to the catalytic activity of PP2A.

**Fig. 7.**
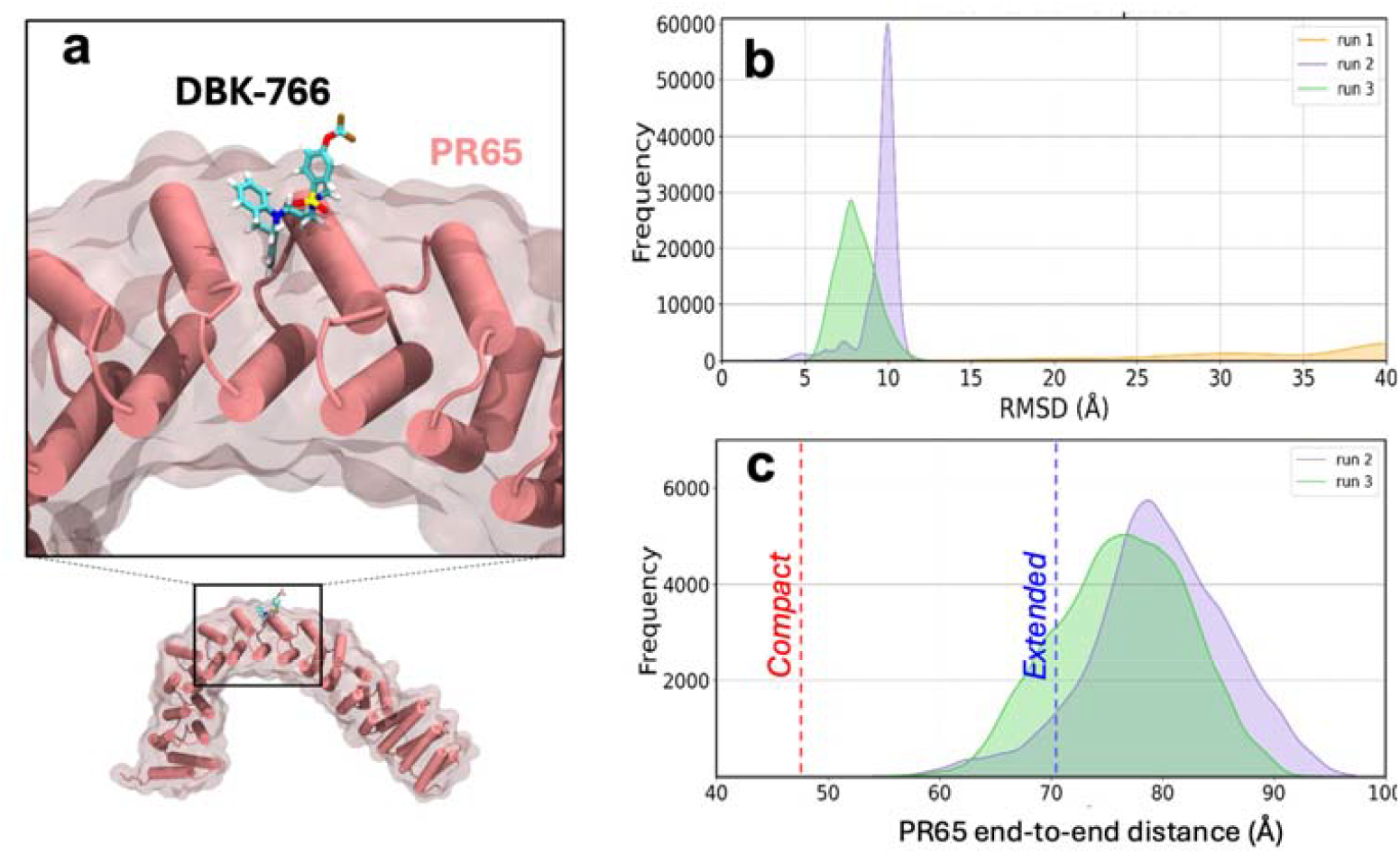
DBK-766 binding at site S_3_ of PR65, coordinated by the outer helices of repeat units 7 and 8. **a)** DBK-766-bound PR65 predicted by computational docking simulations and verified by MD simulations to be a stable pose in two of three runs initiated with the bound form. **b)** RMSD distribution of DBK-766 when bound to that new site, calculated from MD snapshots shows stable binding in two runs (*green* and *purple*) but dissociation in the third (*orange*). **c)** Distribution of PR65’s end-to-end distance deduced from MD snapshots. Results are shown for the two runs that maintained DBK-766 in the bound state.

## Discussion

Repeat-protein folding has been widely studied^27–30,37–39^, and their properties have been shown to be distinct from those of globular proteins, reflecting the linear, repetitive, non-globular architecture. Here we explore the mechanism of action of small molecule activators of PP2A (SMAPs)^10–15^ on the dynamics of HEAT-repeat protein PR65. The mechanics of repeats proteins are much less well studied than those of globular proteins. Helical repeat proteins like PR65 are mechanically weak, and consequently optical tweezer are better than atomic force microscopy for exploring their low-force regime^40–43^. We find that PR65 unfolding under force is very heterogeneous, which is reminiscent of the chemical-induced unfolding pathways of repeat proteins arising due to their structural symmetry. Strikingly, ATUX-8385 binding has a global stabilizing effect on PR65 and prevents misfolding events. This behaviour is similar to molecular chaperones such as the heat-shock proteins and has been mimicked by small-molecule “pharmacological” chaperones^20,44,45^ that have been developed as drugs to prevent proteins from unfolding and misfolding and restore stability to destabilising mutations associated with diseases such as cancer, cystic fibrosis, and lysosomal storage disorders by stabilising the native state of PR65 and aiding its folding. In contrast to ATUX-83853, the non-functional SMAP has a very different mode of interaction and following the first response to stretch-relax cycle (which indicates binding albeit weaker than ATUX-8385), it appears to lack local structure formation/dissolution steps and instead exhibits a smooth response indicative of unfolded/disordered state and absence of PR65 refolding.

The distinctive behavior of the two SMAPs, ATUX-8385 and DBK-766, could be traced back to their different PR65-binding propensities, confirmed by both docking and extensive MD simulations. In this respect, the site S3 emerged as a major hot spot for ATUX-8385 binding. S3 is distinguished by the properties: (i) Its close proximity to the site S_exp_ resolved by cryo-EM for another SMAP, DT-061; S3 and S_exp_ share coordinating residues such as D105. This suggests that S3 may serve as an intermediate site accessible to PR65 prior to its complexation with the regulatory and catalytic subunits of PP2A; trimerization presumably induces a rearrangement to optimize the interactions of the SMAP with all three subunits as resolved by cryo-EM; (ii) ability to promote the open form of PR65, in favor of accommodating the insertion of the two other subunits; (iii) adaptability to alternative poses of the SMAP (including an overall conformational flip) noted in **Fig. 5** and **Fig. S5**, which further attests to the predisposition to induced fit as well as to the contribution of entropic effects to the selection of S3 by ATUX-8385. These three properties support the binding of ATUX-8385 to S3 and its function as an activator. Furthermore, the computed binding affinity of ATUX-8385 to S3 was comparable to the experimental *K*_*D*_ values.

In contrast, DBK-766 preferentially binds a site on the exterior of the repeat units (**Fig. 7** and **Fig. S6**). Even though it also maintains the open position, it would have no effect on the association of PR65 with the subunits B and C, or the stabilization of the active state of PP2A, which explains its lack of functionality as a SMAP. As a final test, we examined whether S3 would bind DT-061 as well, which would strongly support the above inferred hypothesis of its role as a first binding site on PR65 prior to its assembly with the other two subunits. Blind docking simulations unambiguously demonstrate that the site S3 is selected as the highest affinity site by DT-061 (**Fig. S7**) in strong support of the significance of site S3 for (functional) SMAP binding to PR65, before trimerization.

Our previous work suggested that, rather than providing a rigid scaffold, PR65 serves as an elastic connector whose well-defined global motions robustly coordinate cycles of catalysis of multiply phosphorylated substrates by PP2A. Here using single-molecule force microscopy, we probed the mechanical properties of the PR65 ‘nanospring’ and effects of SMAPs, demonstrating that ATUX-8385 acts to modulate the holoenzyme activity by predisposing the PR65 structure to complexation with subunits B and C and by consolidating its intrinsic dynamics, whereas the non-functional DBK-766 disrupts the global dynamics and cooperativity required for PR65 function.

## Materials and Methods

### Expression and purification of PR65

The expression and purification of WT and ybbr-tagged PR65 was performed as previously described ^6^. In brief, plasmid encoding PR65 was transformed into chemically competent C41 *E. coli* (Komander laboratory, MRC-LMB, Cambridge). Cultures were grown at 37°C in 2xYT medium containing ampicillin (50 μg/ml) until an OD_600_ of 0.6 to 0.8 was reached. Protein expression was induced with 500 μM isopropyl-β-d-thiogalactopyranoside (IPTG) (Generon) at 25°C overnight. Cells were harvested by centrifugation at 4000g for 10 min at 4°C before resuspending in lysis buffer (50 mM Tris-HCl pH 7.5, 500 mM NaCl, 2 mM dithiothreitol (DTT)) supplemented with EDTA-free protease inhibitor cocktail (Sigma-Aldrich) and deoxyribonuclease (DNase) I (Sigma-Aldrich). The cells were lysed by passing the suspension two to three times through an Emulsiflex-C5 (AVESTIN) at pressures of 10,000 to 15,000 psi. Soluble protein was separated from cell debris and other insoluble fractions by centrifugation at 35,000g for 35 min at 4°C. The soluble protein fraction was applied to glutathione resin [Amintra Affinity, EGTA (0.5 g/liter), 2 mM DTT], the GST tag was cleaved with thrombin, and PR65 was eluted using a gravity column. After washing the column, the protein was subsequently eluted using a 20× column-volume salt gradient from 0 to 1 M NaCl. MonoQ fractions containing the protein were concentrated before application to a HiLoad 26/600 Superdex 200 pg (GE Healthcare) equilibrated in phosphate-buffered saline (pH 7.4) and 2 mM DTT. Samples were analyzed by SDS-PAGE (polyacrylamide gel electrophoresis) comparing the lysed, flowthrough, and eluted fractions. The identity of the protein was confirmed via MALDI mass spectrometry (Department of Chemistry, University of Cambridge).

### Nano-Differential Scanning Calorimetry (nanoDSF)

PR65 samples (2 µM) in PBS, 2 mM DTT were mixed with ATUX-8385, or with DBK-766, to a final concentration of 100 µM, 10% DMSO and in a total volume of 20 µL. The mixtures were allowed to equilibrate at room temperature for 10 minutes. High-sensitivity capillaries (NanoTemper) were filled with the equilibrated PR65-SMAP mixtures using capillary forces, ensuring no air bubbles were present. The capillaries were loaded into the Prometheus NT.48, and a thermal ramp from 20°C to 90°C at a rate of 1°C per minute was applied. Intrinsic tryptophan and tyrosine fluorescence at 330 nm and 350 nm was continuously monitored. The melting temperature (T_m_) was determined from the first derivative of the fluorescence ratio (I_350_/I_330_) with respect to temperature. The T_m_ corresponds to the inflection point of the melting curve.

### Fluorescence Polarisation (FP)

A fixed concentration of ATUX-8385 (2.5 µM, 5 µM, and 10 µM) was mixed with varying concentrations of PR65 (0 to 80 µM), in PBS + 2 mM DTT, to a total volume of 20 µL in each well of a 384-well microplate (Greiner), and to a final DMSO concentration of 10%. The plate was incubated at 40°C for 30 minutes to ensure binding equilibrium. Fluorescence polarisation was measured at 40°C using an excitation wavelength of 295 ± 10 nm and an emission wavelength of 360 ± 20 nm, on a CLARIOStar plate reader (BMG Labtech), measuring the light in parallel and perpendicular planes relative to the excitation plane. Wells containing only PR65 were used as controls to determine background fluorescence. Fluorescence polarisation (P) was calculated using the equation:

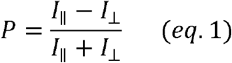

Where *I*_‖_, *I*_⊥_, are the intensities of fluorescence parallel and perpendicular to the excitation plane, respectively.

Non-linear regression analysis was performed using a one-site binding model to fit the data and determine the dissociation constant *k*_*D*_, using the following equation:

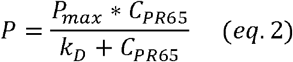

Where *P* is the polarisation, *P*_*max*_ is the maximum polarisation at the plateau, *C*_*PR*65_ is the concentration of PR65 (µM) and *k*_*D*_ is the equilibrium dissociation constant.

### Protein-Ligand Docking Simulations

Docking simulations were performed using AutoDock Vina^46^. In preparation for docking simulations, hydrogens were added to both the receptors and the SMAP, and the resulting structures were saved in PDBQT file format compatible with Vina, utilizing AutoDock Tools1.5.6^47^.Subsequently, partial charges for SMAP were calculated using the Gasteiger method. Simulations were repeated with a series of three-dimensional cubical boxes, with edge lengths ranging from 20 Å to 80 Å and with a grid spacing of 1 Å, centered around K194-L198.

### MD Simulations

In preparation for MD simulations in explicit solvent, each protein-ligand complex was solvated in a truncated octahedral water box using the TIP3P water model, with a minimum distance of 26.0 Å between the solute and the box edge. Na^+^ and Cl^-^ ions were added to balance the charges. All the preparatory steps were conducted using the program *tLEaP*. Three independent MD runs of 800 ns were performed using AMBER20^48^, with the ff14SB^49^ force field for the protein, TIP3P^50^ for the solvent, and GAFF^51^ for the small molecules. Ligand mol2 files were processed with *antechamber*^*52*^ and *parmchk2* to compute AM1-BCC charges^53^. Amber coordinate and topology files were generated using *tLEaP* from AmberTools, upon combining processed ligand mol2 file with apo protein structure. A multi-step protocol was adopted for minimizing and equilibrating each complex, consisting of unrestrained minimization of 2000 steps using the steepest descent method during the first 500 steps to handle large forces, and the conjugate gradient method during the succeeding 1500 steps. A cut-off distance of 10 Å for non-bonded interactions was used. The system underwent 20 ps of restrained NVT equilibration at 298 K, with a 2-fs time step, using a Langevin thermostat with a damping coefficient of 1 ps^−1^ and harmonic restraints (k = 1 kcal/mol/Å^2^) applied to all non-hydrogen atoms except water molecules and ions. This was followed by a 1 ns restrained NPT equilibration at 298 K, using the same Langevin thermostat and the Monte Carlo barostat, with harmonic restraints. A further 1 ns unrestrained NPT equilibration was then performed using the same Langevin thermostat and MC barostat, during which harmonic restraints were removed. The production runs consisted of 800 ns of NPT simulations at 298 K with a 2 fs time step, using the Langevin thermostat and MC barostat to maintain pressure at 1 atm. Non-bonded interactions were calculated with a 10 Å cut-off, and long-range electrostatics were treated using the particle-mesh Ewald method, with trajectory data saved every 10 ps, and viewed using PyMol and VMD. Trajectory analysis was performed using the *cpptraj* ^*54*^ program.

### Protein-DNA constructs

PR65 with ybbR tags at the N- and C-termini was conjugated to CoA modified DNA oligos (biomers) in a 50 µl Sfp-synthase reaction buffer using 10 µM protein, 10 µM Sfp-synthase and 20 µM CoA oligos. The reaction was kept for overnight at RT. Next the reaction mixture was purified by size exclusion chromatography using a Superdex S200 10-300 equilibrated in 50 mM Tris-HCl, 150 mM NaCl, 1mM DTT ^22^.

DNA handles of size 600 bp were PCR amplified from λ-DNA (Jena Bioscience) using a triple biotinylated primer, a triple digoxigenin modified primer and a primer with abasic site (biomers). A standard PCR was performed using Q5 master mix DNA polymerase (NEB) at an annealing temperature of 60ºC and elongation temperature of 68ºC. This PCR produces DNA handles with 5’-overhangs complementary to the CoA modified oligos. The DNA handles were next purified from agarose gel extraction followed by QIAquick PCR purification protocol. Different dilutions of oligos conjugated PR65 was mixed with 200 ng DNA handles, and the reaction was checked by 1% agarose gel to find the optimum concentration for the DNA-oligos hybridization. Before optical tweezers measurements, the samples were prepared from fresh hybridization reaction on the same day.

### Optical tweezers experiments

Streptavidin and anti-digoxigenin coated beads (2 µm) were purchased from Spherotech and stored at 4°C until use. 10 ng of PR65 constructs were mixed with 1 μl anti-digoxigenin beads in 10 μl Tris-HCl buffer (50 mM Tris-HCl pH 7.5, 150 mM NaCl, 2 mM DTT). The reaction mixture was then incubated at 4 °C for 30 min. Next, the protein coated anti-digoxigenin beads were dissolved in 450 μl Tris-HCl buffer for optical tweezers experiments. Optical tweezers measurements for without the SMAP condition (**Fig.1**) were done in Tris-HCl buffer with 2% DMSO. For measurements done in presence of SMAP, 10 μM SMAP were dissolved in 2 % DMSO and Tris-HCl buffer. Constant velocity experiments at 100nm/s were performed on a dual trap optical tweezers instrument (C-trap from Lumicks). Tethers were formed by bringing PR65 construct-coated and Streptavidin beads in proximity. The protein was stretched and relaxed by moving one of the traps. The data was acquired at 78 kHz and averaged to 100 Hz.

## Data analysis

To confirm that the data corresponded to a single tether, the following checks were performed: the total measured unfolded length was compared with the expected length and tether breakage in a single rupture event. The quantification of unfolding forces, contour lengths and refolding forces were done from force extension data, using a custom-built Python code. Unfolding and refolding forces of each transition were determined as the average force before a transition of minimum 10 nm size. The absolute contour lengths (L_c_) were quantified from Worm-like chain (WLC) fitting of the force extension curves. The persistence lengths of the DNA (30 nm) and protein (0.5 nm), the stretch modulus of DNA (400-600 pN) and the protein (300 pN) were fixed parameters in the WLC model.

## Statistical analysis

The statistical significance of differences in unfolding and refolding force distributions between experimental conditions was calculated using two samples assuming unequal variance t-Test. Test results are mentioned as *p* values in the main text. In box charts, whiskers indicate 90% and 10% extreme values, the inner line represents the median, the length of the box indicates interquartile range and the inner small square the mean of the population.

## Data availability

All data supporting the findings of this study are available in the main text and Supporting Information figures. The raw data that support the findings of this study are available from the corresponding authors upon reasonable request.

## Code availability

The Python code used for analysis is available from the corresponding author upon reasonable request.

## Acknowledgments

LI, IB, S-HY, and RG gratefully acknowledge support from HFSP grant RGP0027/2020 and LI from BBSRC (Biotechnology and Biological Sciences Research Council) grant BB/T002697/1. Support from National Institutes of Health awards R01 GM139297 and R01 DK116780 is gratefully acknowledged by IB. We thank Dr Marie Synakewicz for helpful advice on protein purification and sample preparation.

## Conflict of Interest

The authors have no conflicts of interest.

## Notes

### Competing Interest Statement

The authors have declared no competing interest.

