## Supporting Information for "Small Molecule Activators of Protein Phosphatase 2A Exert Global Stabilising Effects on the Scaffold PR65"

**`**

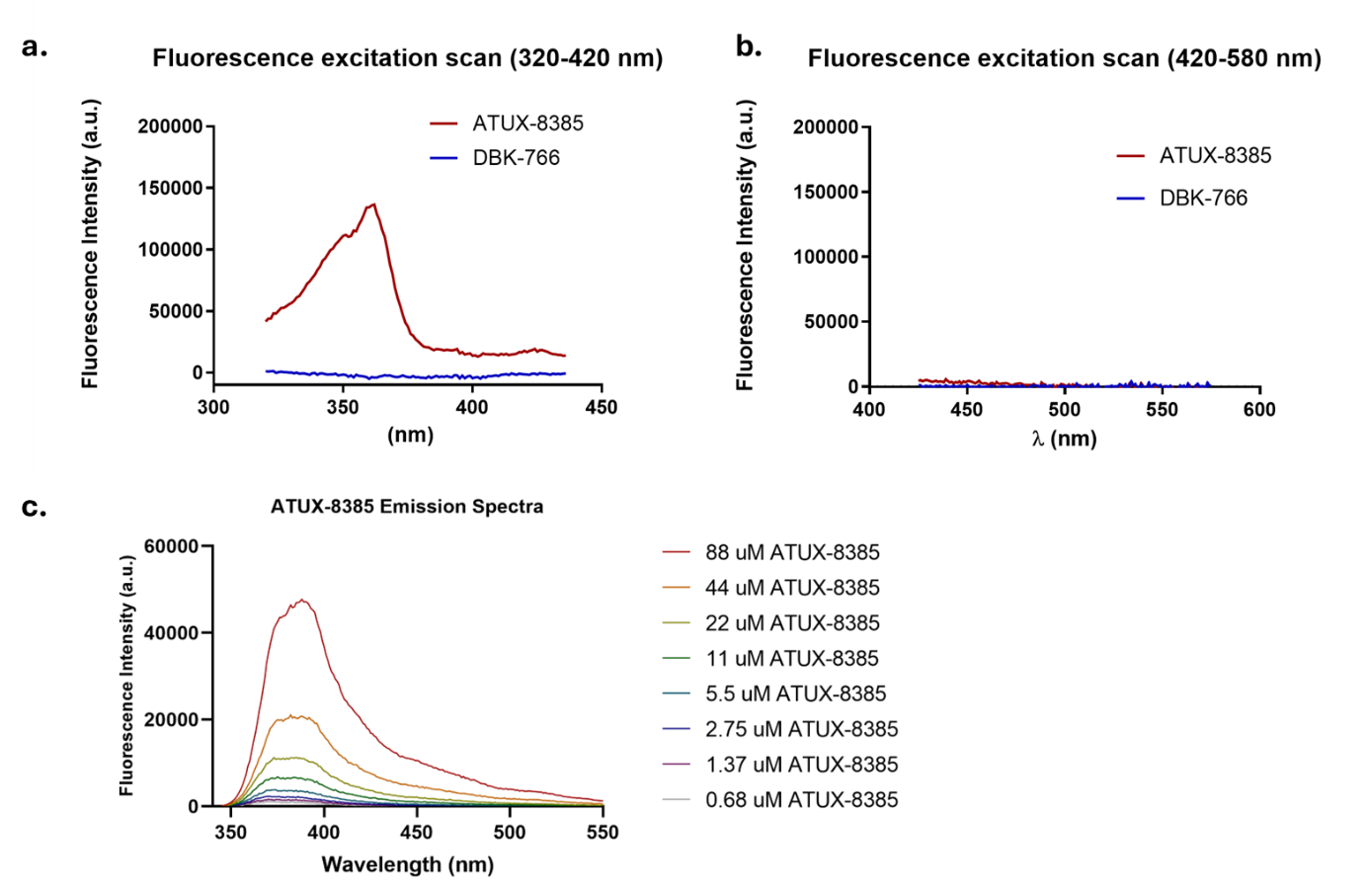

**Fig. S1: Fluorescence spectra of ATUX-8385 and DBK-766.** Fluorescence excitation scans from 320-420 nm **(a)** and 420-580 nm **(b)** of ATUX-8385 and DBK-766 show that ATUX-8385 has an excitation maximum at ~350 nm, whereas DBK-766 is not fluorescent. **c)** Fluorescence emission spectrum of ATUX-8385 upon excitation at 290 nm.

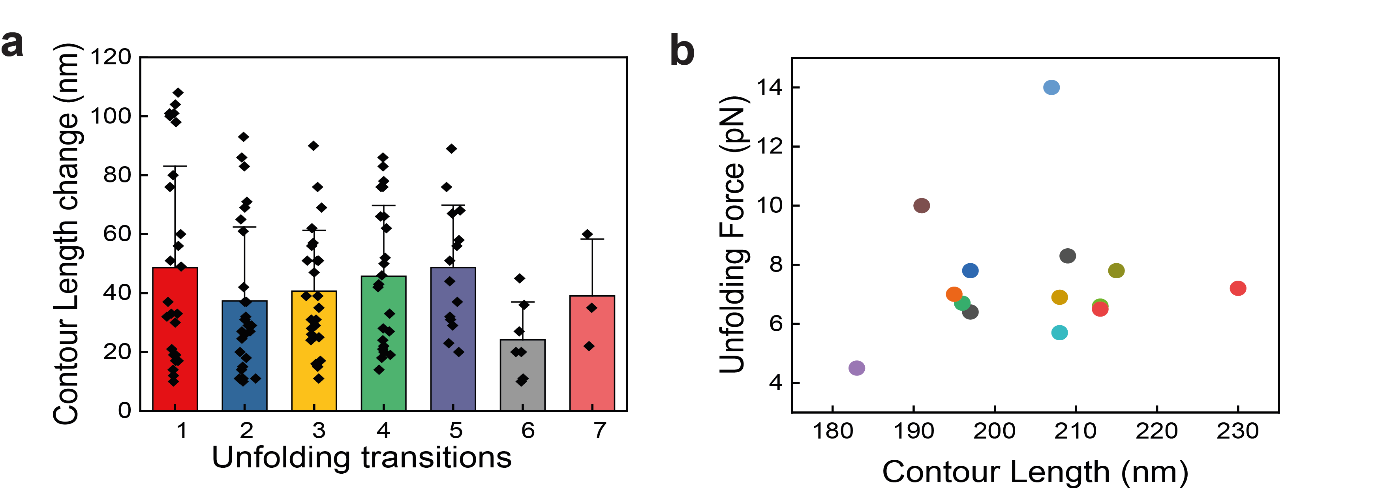

**Fig. S2. Contour length variations and dependency on unfolding force observed in experiments. a)** Distribution of contour length changes associated with individual unfolding transitions measured from all the stretching curves of PR65 in the absence of SMAP. **b)** Maximum unfolding force vs absolute contour length of the fully unfolded molecule measured from the first pulls of all the PR65 molecules in the absence of SMAP.

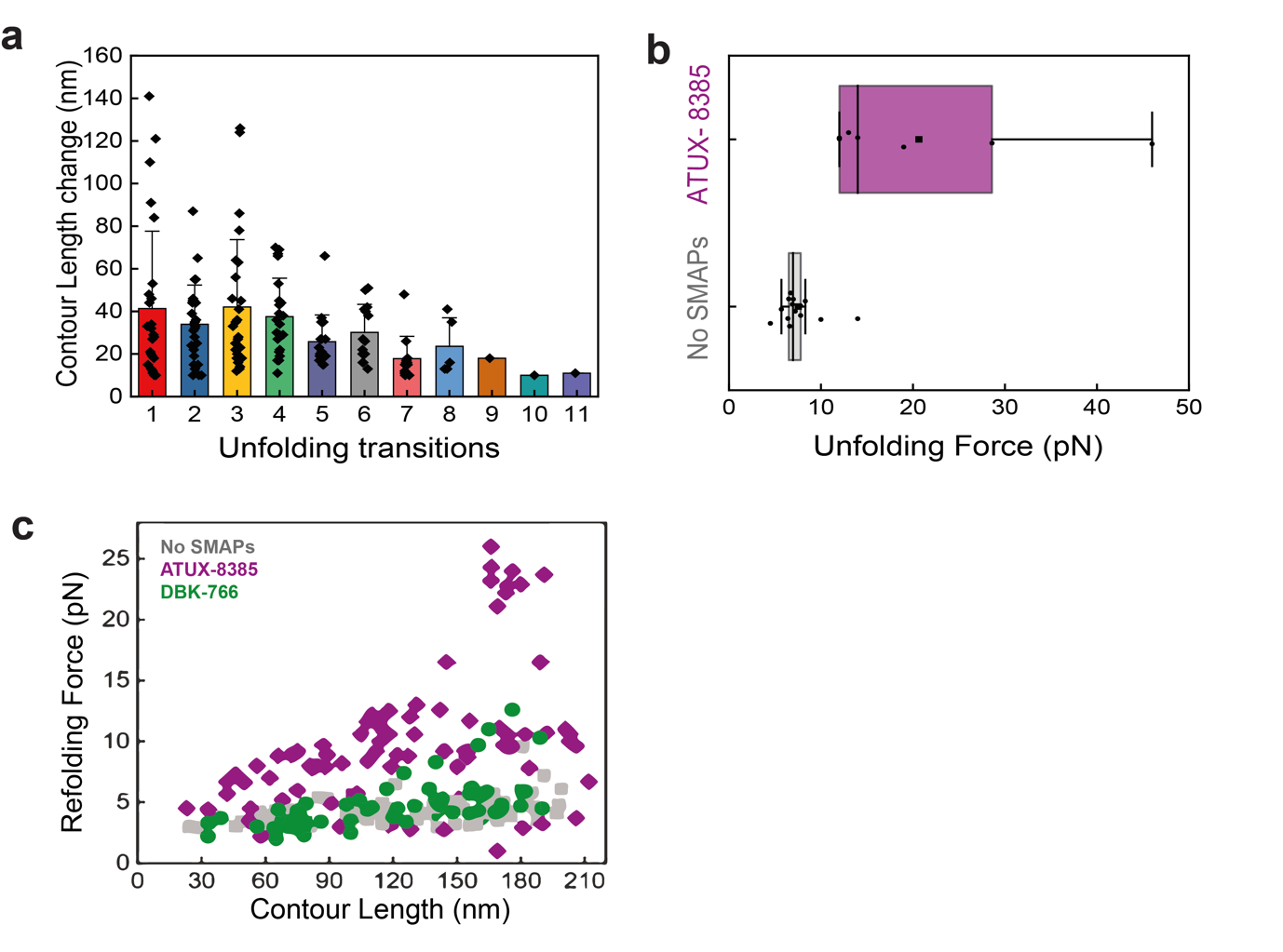

**Fig. S3. a)** Distribution of contour length changes associated with each unfolding transition measured from all the stretching curves of PR65 in the presence of ATUX-8385. **b)** Box plots showing distribution of maximum unfolding force of the first pulls for PR65 with (*purple*) and without (*gray*) SMAP. **(c)** Refolding force *vs* absolute contour length (L_c_) of each intermediate state observed during relaxation of apo PR65 (*gray squares*), and PR65 bound to ATUX-8385 (*purple diamonds*) and DBK-766 (*green circles*).

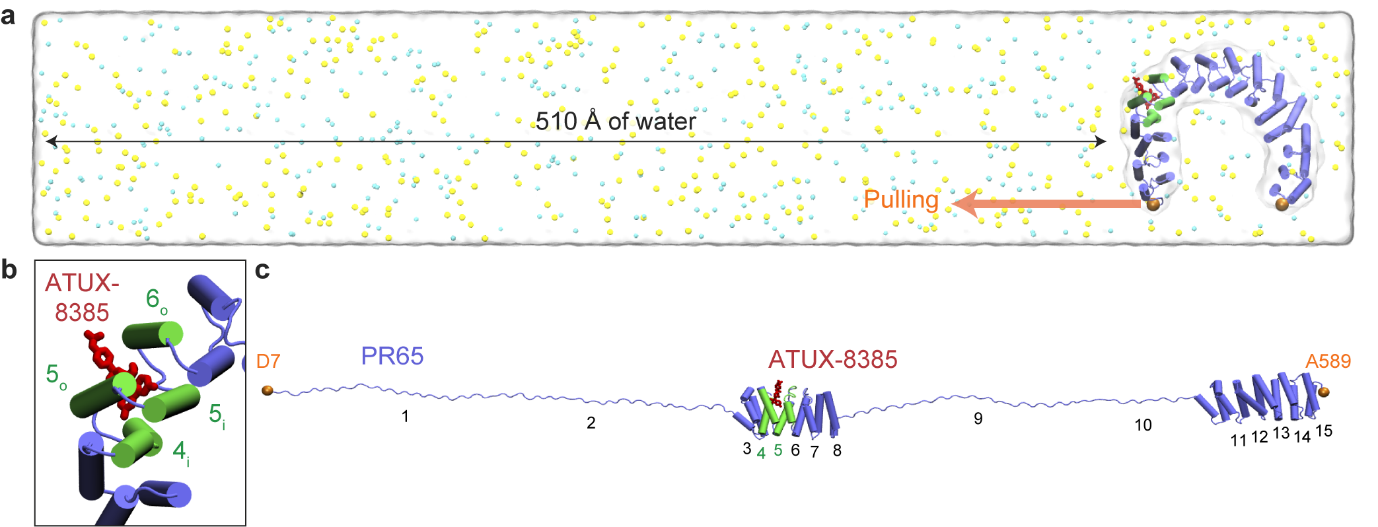

**Fig. S4. SMD simulations of PR65 bound to ATUX-8385. a)** PR65 bound to ATUX-8385 solvated in a water box. Helices interacting directly with ATUX-8385 are highlighted in *green*, and non-interacting helices are colored *blue*. *Cyan* and *yellow* beads in solution represent Na⁺ and Cl⁻ ions, respectively. The steered and fixed atoms are shown with *orange beads*. **b)** A magnified view of the ATUX-8385 binding site based on docking simulations onto the compact conformation of PR65. **c)** The conformation of PR65 after completion of the first SMD simulation run with ATUX-8385 bound, following application of a pulling force over a distance of 50 nm.

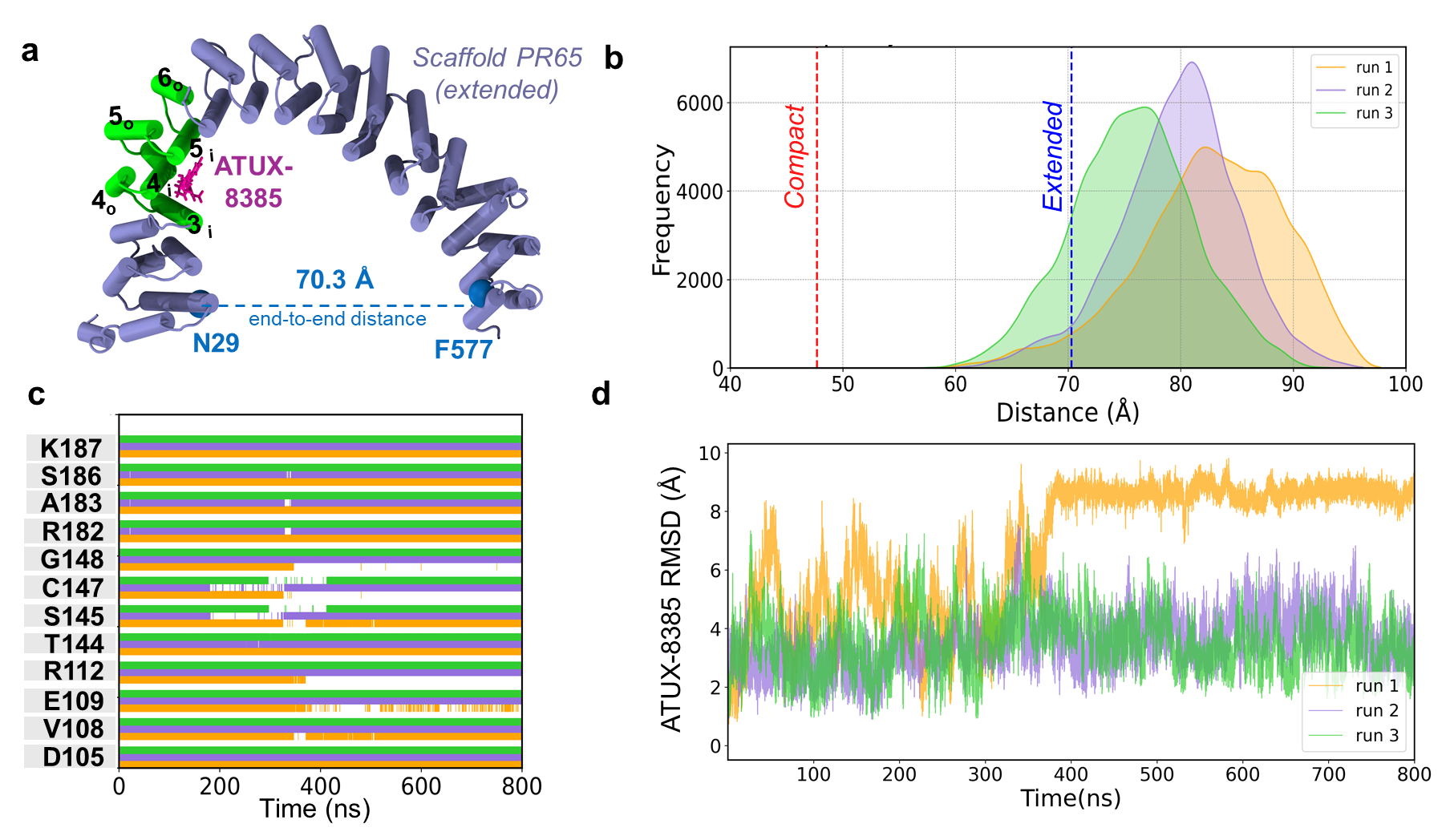

**Fig. S5. Stability of ATUX-8385–bound PR65 examined by MD simulations.** **a)** Binding pose of ATUX-8385 to PR65. The diagram shows the initial conformation used in MD simulations. **b)** Histogram of end-to-end distances sampled during MD simulations. The histograms for the three runs are shown in different colors. **c)** Time evolution of ATUX-8385 binding with coordinating PR65 residues. Coordinating residues are those making atom-atom contacts within a cutoff distance of 5 Å across three independent runs, represented by *orange*, *purple*, and *green*, respectively. Blank regions refer to cases where those particular residues were more than 5 Å away from ATUX-8385. **d)** ATUX-8385 RMSD from MD snapshots. In run1, ATUX-8385 undergoes a rotational motion at about 400 ns and remains bound at the same site, but with a different orientation.

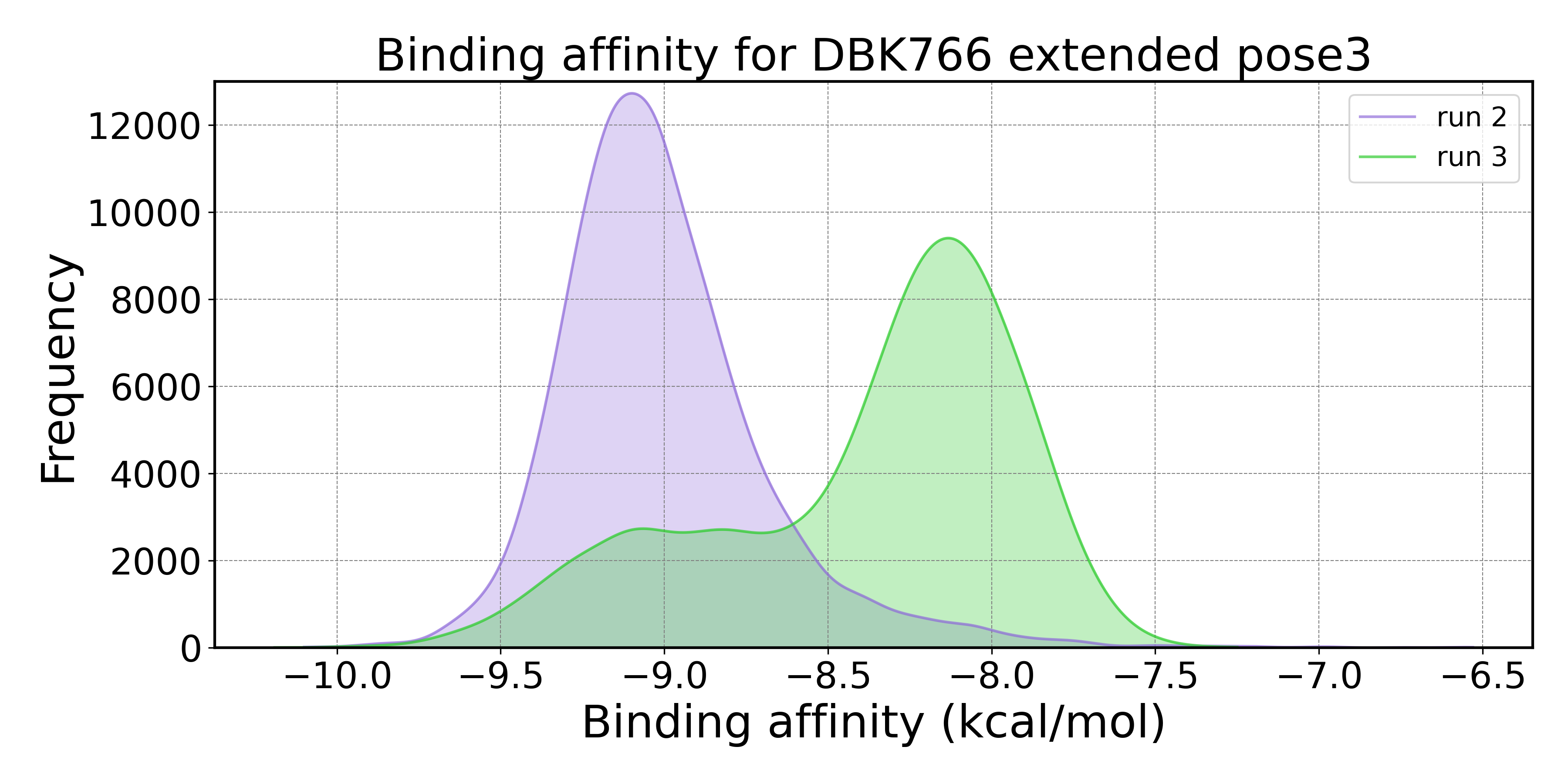

**Fig. S6. Distribution of DBK-766 binding energies observed for DBK-766 bound to S_3_.** Results from two runs are shown, in which DBK-766 remained bound to PR65.

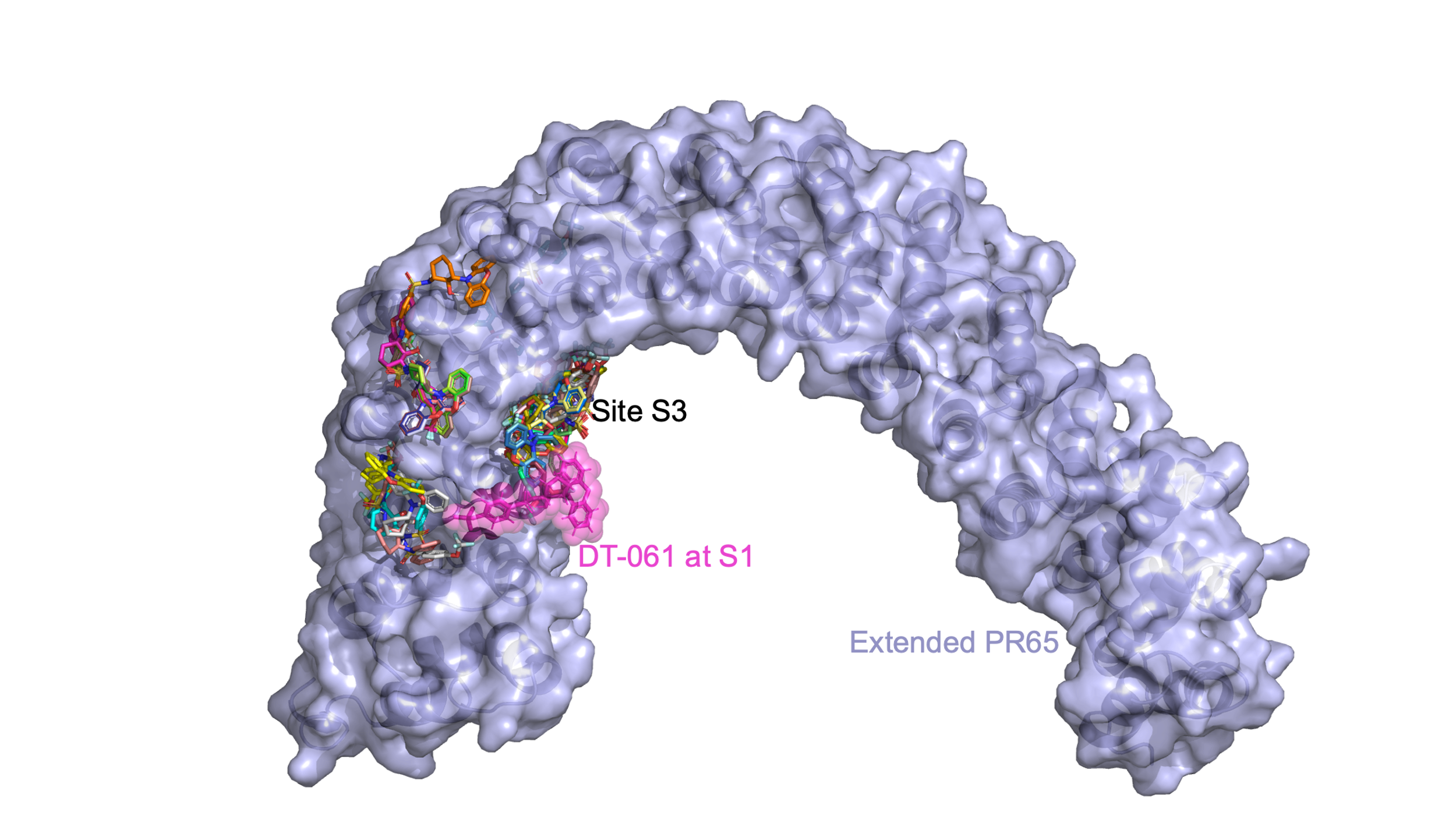

**Fig. S7. Results from docking simulations of DT-061 onto extended PR65. Multiple binding poses are shown (multicolor sticks).** Site S3 is distinguished by its high affinity to bind DT-061 in multiple runs. For comparison DT-061 *(pink sticks and shade*) is also shown at the site (S1) resolved by cryo-EM for the trimeric PP2A. The Vina binding affinity of the best docked pose at S3 is -7.88 kcal/mol, and that of PRODIGY-LIG is -9.48 kcal/mol.
